# Ganglioside profiling by ion mobility and mass spectrometry imaging identifies distinct biomolecular isomers in Niemann-Pick Disease

**DOI:** 10.64898/2026.09.17.752326

**Authors:** Signe Frost Frederiksen, Pelayo A. Penanes, Charlotte Laurfelt Munch Rasmussen, Annette Burkhart, Christian W. Heegaard, Daniel Wüstner, Ole N. Jensen

**Author notes:** **Corresponding Author: Prof. Ole N. Jensen:** Department of Biochemistry and Molecular Biology, University of Southern Denmark, Denmark.

## Abstract

Gangliosides are glycosphingolipids implicated in the neurodegenerative pathology of Niemann-Pick disease type C2 (NPC2 disease), where accumulation of GM2 and GM3 is a hallmark of disrupted lipid trafficking and disease progression. Some gangliosides are structural isomers with identical molecular composition and mass, making them difficult to distinguish by mass spectrometry imaging (MSI) alone and leaving their spatial distributions unresolved. To address this challenge, we combined ion mobility spectrometry with mass spectrometry imaging for isomer-resolved ganglioside analysis.

We assessed ganglioside complexity in wild-type (*Npc2+/+*) and NPC2-deficient mice with or without AAV-BR1-mediated NPC2 gene therapy (*Npc2-/-* AAV-BR1-NPC2 and *Npc2-/-* vehicle) using matrix-assisted laser desorption ionization (MALDI) MSI, and imaging-parallel reaction monitoring-parallel accumulation serial fragmentation (iPRM-PASEF-MS/MS) for *in situ* brain tissue analysis. Trapped ion mobility spectrometry (TIMS) enabled gas-phase separation of isomeric gangliosides, while iPRM-PASEF-MS/MS facilitated structural discrimination based on diagnostic fragment ions and identification of molecular modifications. Brain sections were analyzed by MALDI-MSI at 20μm spatial resolution, with selected regions examined using post-ionization imaging by MALDI-2 at 5μm. 59 mobility-resolved ganglioside features representing 23 putative annotations were identified, including separation and spatial mapping of GM1a/GM1b and GD1a/GD1b isomers. Notably, unreported modifications of gangliosides were identified, including GM1 fucosylation in *Npc2*-/- mice and GD1 O-acetylation exclusively detected in *Npc2*+/+ mice. Spatial ganglioside profiling by MSI demonstrated that gene therapy of *Npc2*-/- mice partially restored normal ganglioside localization in brain, indicating modulation of lipid storage. This study establishes isomer-resolved lipid imaging for investigating ganglioside alterations in disease models and the effects of therapeutic intervention.

---

Niemann-Pick disease type C (NPC disease) is an autosomal recessive lysosomal storage disorder (LSD) caused by loss of function of either NPC1, a lysosomal membrane protein binding cholesterol, oxysterols, and sphingosine; or of the soluble lysosomal cholesterol-binding protein NPC2. Deficiency of NPC1 and NPC2 impair cholesterol trafficking and drives secondary accumulation of substrates, such as glycosphingolipids, within the lysosomal compartments [1–3]. As for many LSDs, these metabolic defects lead to progressive cellular dysfunction that may culminate in premature death, partly driven by neurodegeneration [4, 5]. In brain tissue, this includes extensive lysosomal accumulation of gangliosides (GGs), such as GM1, GM2, GM3, and GD1, contributing to disease progression [1].

GGs are sialylated glycosphingolipids enriched in neuronal membranes, where they regulate synaptic stability, signal transduction, and immune interactions [6, 7]. Their structural diversity arises from variations in both the oligosaccharide headgroup and the ceramide backbone. These variations generate numerous isomeric species that share the same molecular composition but differ in the structural arrangement of monosaccharide residues within the oligosaccharide chain. This is exemplified by GM1a and GM1b, which have the same molecular composition but differ in the position of their single sialic acid residue within the oligosaccharide headgroup, resulting in distinct isomeric structures with either terminal or internal sialic acid arrangement. Structural variation of the ceramide moiety, including chemical configuration, double-bond positions, and carbon-chain length, further contributes to isobaric and isomeric GG species. Sialic acid residues attached to the oligosaccharide chain can further be subjected to O-acetylation (O-Ac) at the C4-, C-7, C-8, or C-9 hydroxyl groups, generating derivatives with distinct biochemical properties [8, 9]. Furthermore, additional monosaccharides, such as fucose, can be conjugated to the carbohydrate moiety of GGs [10]. Because of this extensive structural diversity arising from variations in glycan composition, chemical modifications of the carbohydrate residues, ceramide structure, and the presence of isomeric species, comprehensive structural and functional characterization of GGs presents a major analytical challenge [11].

Whereas GG accumulation is well described, particularly for NPC1 disease, most studies reported bulk biochemical or immunohistochemistry analyses that did not resolve structural GG isomer-level identification together with their spatial distributions within the mammalian brain [2, 3, 12]. This distinction is important, as structurally related GG isoforms may exhibit differences in biological function, metabolism, and regional localization, reflecting impaired lipid trafficking in NPC disease and variations in cellular composition. Consequently, it remains unclear whether specific GG isomers display region-dependent distributions and how such difference may contribute to disease pathology.

Matrix-assisted laser desorption/ionization mass spectrometry imaging (MALDI-MSI) provides a probe- and label-free approach for detecting, identifying, and visualizing the spatial distribution of multiple molecular species directly in tissue [13]. MALDI-MSI is particularly well suited for investigating region-specific lipid alterations associated with LSDs. However, most MALDI-MSI studies of GGs rely solely on biomolecular identification by exact mass determination (MS^1^-level analysis and annotation), which is insufficient to distinguish structural isomers with identical elemental composition and identical molecular mass [14].

Trapped ion mobility spectrometry (TIMS) adds an orthogonal separation dimension by resolving ions according to differences in their gas-phase mobility as they move through a buffer gas against an opposing electric field. Measured ion mobility is converted into ion-specific collisional cross section (CCS) values, which provide information on differences in gas-phase conformation and enable separation of ions with identical molecular compositions, but structurally different, as observed with gangliosides [15].

TIMS-MSI studies have demonstrated the potential to visualize region-specific GG isomer distributions in rodent tissue, including the separation of structurally similar gangliosides isomers such as GD1a and GD1b, and multiple GM1 variants, based on their distinct mobilities [11, 16, 17]. Nevertheless, mobility values alone cannot offer definitive structural assignments, and without additional structural elucidation, the identification of GG species remains tentative [15]. TIMS-MSI studies have therefore relied on targeted tandem mass spectrometry (MS/MS) analysis of individual precursor ions for structural confirmation using diagnostic fragment ions, including losses of hexoses, sialic acids, and neutral losses of carbon dioxide, and water. [17–19]. Although informative, a single precursor-targeted approach restricts the number of isomers that can be structurally characterized within a single imaging experiment. Imaging-parallel reaction monitoring-parallel accumulation serial fragmentation-MS/MS (iPRM-PASEF-MS/MS) tackles these drawbacks by enabling detailed MS/MS analysis of multiple precursor ions during a full imaging run. This approach allows for parallel acquisition of high-quality MS/MS data *in situ* from tissue, including analysis of gangliosides and their isomers while preserving their spatial context [20, 21].

In this study, we applied ion mobility spectrometry in combination with MS imaging to detect known and previously unresolved ganglioside species in normal and perturbed brain tissues, including isobaric and isomeric GGs, in the context of Nieman-Pick Disease. Using MALDI-TIMS MSI in combination with iPRM-PASEF-MS/MS, we determined the spatial distributions and molecular heterogeneity of cerebral gangliosides in brain sections from wild-type mice (*Npc2+/+*), and in age-matched NPC2-deficient littermates subjected to either AAV-BR1-mediated NPC2 gene therapy (Npc2-/- AAV-BR1-NPC2) or vehicle controls (*Npc2-/- vehicle).* The objectives were to (i) detect and spatially map an extensive set of gangliosides in context of NPC2 pathology, (ii) differentiate structural variants and GG isomers at the MS^1^-level using high TIMS resolving power, and (iii) structurally elucidate selected isomeric GG species directly in brain tissue. The analytical strategy presented combines ion mobility separation with spatially resolved tandem MS to provide detailed insights into ganglioside structures and their spatial distributions in NPC2 pathology, potentially facilitating the identification of novel molecular targets for therapeutic interventions.

## Experimental Section

### Animal approval and reporting

NPC2-deficient mice were bred and maintained at Aarhus University, Aarhus, Denmark, as previously described [1]. Breeding and all animal procedures were approved by the Danish Animal Experiments Inspectorate (license no. 2018-15-0201-01467 and 2019-15-0202-0056) and complied with the Danish Animal Experimentation Act (BEK no. 1065 of 25/09/2024) and the European Directive 2010/63/EU and adhered to the ARRIVE guidelines. We studied age-matched hemispheres from 16 mouse brains (12 weeks of age); wild-type mice (*Npc2+/+*), *Npc2-/-* mice that received gene therapy with an adeno-associated virus (AAV) carrying the NPC2 gene, specifically targeting brain endothelial (BR1) cells (*Npc2-/-* AAV-BR1-NPC2), and their matching vehicle controls (*Npc2-/-* vehicle) [1, 22].

### Tissue sample preparation for MALDI MSI

Mice were euthanized under deep isoflurane anesthesia by transcardial perfusion with cold PBS. Hemisected mouse brains were wrapped in aluminum foil and rapidly frozen in a solution of 70% ethanol mixed with dry ice. Due to the high lipid content in *Npc2-/-* brains, the tissue was notably soft during initial cryosection tests, which impaired the adherence to Optimal Cutting Temperature (OCT) medium (VWR Chemicals) and increased the risk of tissue damage. A total of four *Npc2+/+,* five *Npc2-/- vehicle*, five *Npc2-/-* AAV-BR1-NPC2, were successfully prepared, as previously reported [1].

### Tissue embedding and cryosectioning

Brain tissue was stabilized to maintain integrity and morphology and to ensure successful sectioning by embedding in 5% (w/v) carboxymethyl cellulose (CMC; molecular weight ∼700.000, Sigma-Aldrich). Brain tissues were subsequently frozen using 70% ethanol (VWR Chemicals) in dry ice and stored at -80°C until further use. The tissue was cut into 10 µm thick sections using a Leica CM1860 cryostat (Leica Biosystems) with temperature set to -25°C. Sections were thaw-mounted onto IntelliSlides (Bruker Daltonics, Bremen, Germany) and stored at -80°C until further use.

### Matrix application for MALDI MSI

Matrix application was performed using a HTX M3+-sprayer (HTX technologies, LLC) with 2.5-dihydroxyacetophenone (DHAP) (Bruker Daltonics, Bremen), dissolved in 70% ethanol with 0.1% trifluoracetic acid (VWR Chemicals). Spraying conditions are provided in Supplementary Table 1.

### Mass spectrometry

#### MALDI-TIMS-MSI Data Acquisition

TIMS-MSI data acquisition was conducted using a timsTOF fleX MALDI-2/MicroGRID mass spectrometer (Bruker Daltonics, Bremen), operated in negative ion mode with TIMS activated. The gas pressure was adjusted to 2.35 mbar to enable a larger mobility range and to capture the GGs. The mass range was set to *m/z* 500-2300, and the mobility window ranged from 1/K_0_ 0.80-2.49 V·s/cm^2^, with a ramp time of 1000ms, corresponding to an accumulation time of 16.9ms. All experiments were carried out in standard MALDI mode with laser repetition rate of 10kHz at 20 µm spatial resolution and 180 laser shots per pixel (Fig. 2A).

#### MALDI iPRM-PASEF-MS/MS Data Acquisition

For iPRM-PASEF MS/MS experiments the instrument mass range was set to *m/z* 50-2500 and all ion transfer settings were adapted to detect fragment ions in this range. For each 40 µm pixels measured during MSI, the analyzed area was divided into four 20µm sub-pixels (quadrants). The first quadrant was used for standard MS acquisition, while the remaining three quadrants were used sequentially for targeted MS/MS acquisition (msms1, msms2, and msms3) using the iPRM-PASEF mode. To reduce the risk of overlapping isolation windows – particularly important for structurally similar ganglioside isomers – precursors were distributed across three separate target list, each used in a different MS/MS acquisition cycle (Supplementary Table 2). Precursors were isolated based on their mobility (1/k_0_) values and fragmented using collision-induced dissociation (CID). Collisional energies of 80eV, 95eV, and 100eV were applied at 1/K_0_ values of 0.80, 1.60, and 2.20 V·s/cm^2^, respectively, with intermediate values determined by linear interpolation. Precursor ions were isolated using a 5 Da window at *m/z* 622 and *m/z* 1521 (Fig. 2B).

#### MALDI MSI with microGRID 5μm

For high spatial resolution (5 μm) MALDI images, the DHAP matrix layer was first applied to the IntelliSlides (Bruker Daltonics, Bremen, Germany) using the HTX M3+ sprayer (HTX technologies, LLC), and the tissue section was placed on top. This setup reduces pixel oversampling and enhances structural definition within the image. Because the laser must penetrate the tissue to reach the underlaying matrix, higher laser power of 50% was used with the consequence of little to no tissue remaining after measurements. MSI data was acquired at a raster size of 5 μm, 5 shots per pixel, using post ionization (MALDI-2) with 5 μs delay.

### Data Processing and Analysis

MSI MS^1^ data was processed using SCiLS lab 2026b (Bruker Daltonics), with each spectrum normalized to its root mean square (RMS) intensity. Feature identification and annotation were conducted through SCiLS lab using Metaboscape 2025b (Bruker Daltonics). Annotations were performed using a Lipidmaps.csv *(Download: 16.01.2025)* target list with the acceptance parameters: mass error 5-10ppm, CCS values within 2-5% of deviation. The parameters were kept the same for the iPRM-PASEF-MS/MS annotation with the addition of the MS/MS score. To ensure comprehensive coverage of ganglioside signals while excluding noise, the MS/MS match score threshold was constrained between 100-600. All MS/MS fragment data were analyzed using Compass DataAnalysis 6.1 (Bruker Daltonics). Fragmentation spectra of the precursor ions were extracted from the mobility heatmaps based on their 1/K_0_ values. De-smoothed mobility plots were evaluated to check whether two mobilities fell within the same window and to avoid overlapping ion mobility distributions. Ganglioside species were identified based on accurate mass determination, CCS value, and MS/MS fragmentation patterns generated *in-situ* from LipidMaps sphingolipids fragmentation tool [23]. The fragmentation tool predicts the oligosaccharide fragments based on the potential Y, B; Z, C nomenclature with the possibility of adding cross-link fragmentation. The ceramide ion fragments are denoted O, S, T, G, U, V, G, E, P, H, depending on the fragment location in the ceramide tail (Supplementary Figure 1) [6].

## Results and Discussion

### Ganglioside profiles reveal distinct patterns and isomer complexity in NPC2-deficient mouse brain

In NPC2 deficiency, altered cholesterol trafficking is accompanied by the accumulation of gangliosides (GGs) associated with neurodegenerative and neuroinflammatory processes in the brain [24]. To investigate these changes at the molecular level, brain sections from *Npc2+/+, Npc2-/-,* and AAV-BR1-NPC2 treated *Npc2-/-* mice were analyzed using MALDI-TIMS-MSI. At the MS^1^-level, differences in GG profiles were prominent between the *Npc2+/+* and *Npc2-/-* samples, reflected by changes in both signal intensity and the distribution of GG species across the *m/z* range (Fig. 1A, C). Focusing on the *m/z* range typically associated with gangliosides (*m/z* 1000-2300) and ion mobility values of 1/K_0_ 1.7-2.5, a total of 63 features defined by their *m/z* and 1/K_0_ values were selected for further analysis (Supplementary Table 2). Of these 63 features, 59 were annotated as GG species by their accurate mass within ±10 ppm error. These 59 annotated features corresponded to 23 unique GG structures, suggesting that numerous GG species were assigned based on the same m/z value despite being resolved in the mobility dimension (Fig. 1B). This likely reflects limitations in the annotation approach, where assignments are primarily based on m/z values, and available CCS prediction tools are not sufficiently optimized for larger lipid species, such as gangliosides. These results and observations indicate that structural information is required to further resolve isomeric ganglioside species.

**Figure 1:**
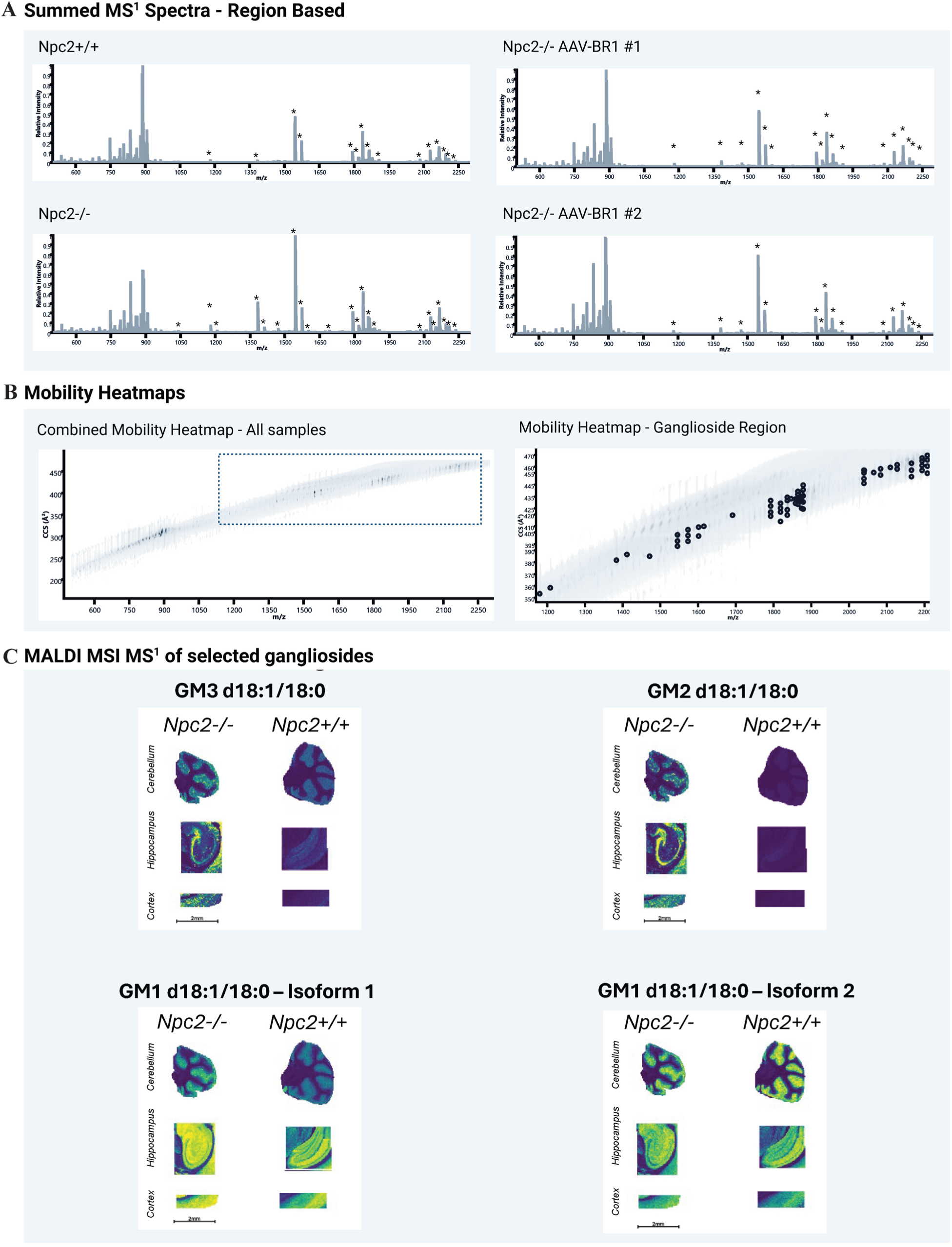
Ganglioside Profiles Reveal Distinct Patterns in NPC2-deficient mouse brains. A) MS^1^-level MALDI-TIMS-MSI spectra shows prominent differences in ganglioside-associated (GG) signals between wild type (Npc2+/+) and NPC2-deficient mice (Npc2-/-), with two representative AAV-BR1-NPC2 treated Npc2-/- samples (#1, #2) displaying intermediate profiles. Peaks corresponding to GG species are indicated with star-icon (*). B) Mobility heatmaps illustrate the distribution of detected ion across the full m/z range, with the ganglioside region (m/z 1000-2300) highlighted. Within this region, multiple mobility-resolved features are observed at similar m/z values, suggesting increased molecular complexity in ganglioside species. C) MSI of a selected panel of gangliosides differentially expressed in the Npc2-/- vs Npc2+/+ mouse model (GM3 d18:1/18:0, GM2 d18:1/18:0, and GM1 d18:1/18:0 Isoform 1) and one ganglioside with similar intensity profile in both models; GM1 d18:1/18:0 Isoform 2. Data were acquired using a timsTOF fleX MALDI-2 microGRID platform in negative ion mode, with a mass range of m/z 500-2300 and mobility range of 0.80-2.49 1/K0, and a TIMS ramp time of 1s.

### Resolving ganglioside isomer complexity by targeted iPRM-PASEF-MS/MS

The detection of multiple mobility-separated features at identical m/z values suggested structural complexity within the detected GG ion species. To determine whether these features represented GG isomers, we employed a targeted iPRM-PASEF-MS/MS workflow, enabling fragmentation of mobility-resolved precursor ions and identification based on diagnostic patterns (Fig. 2).

**Figure 2:**
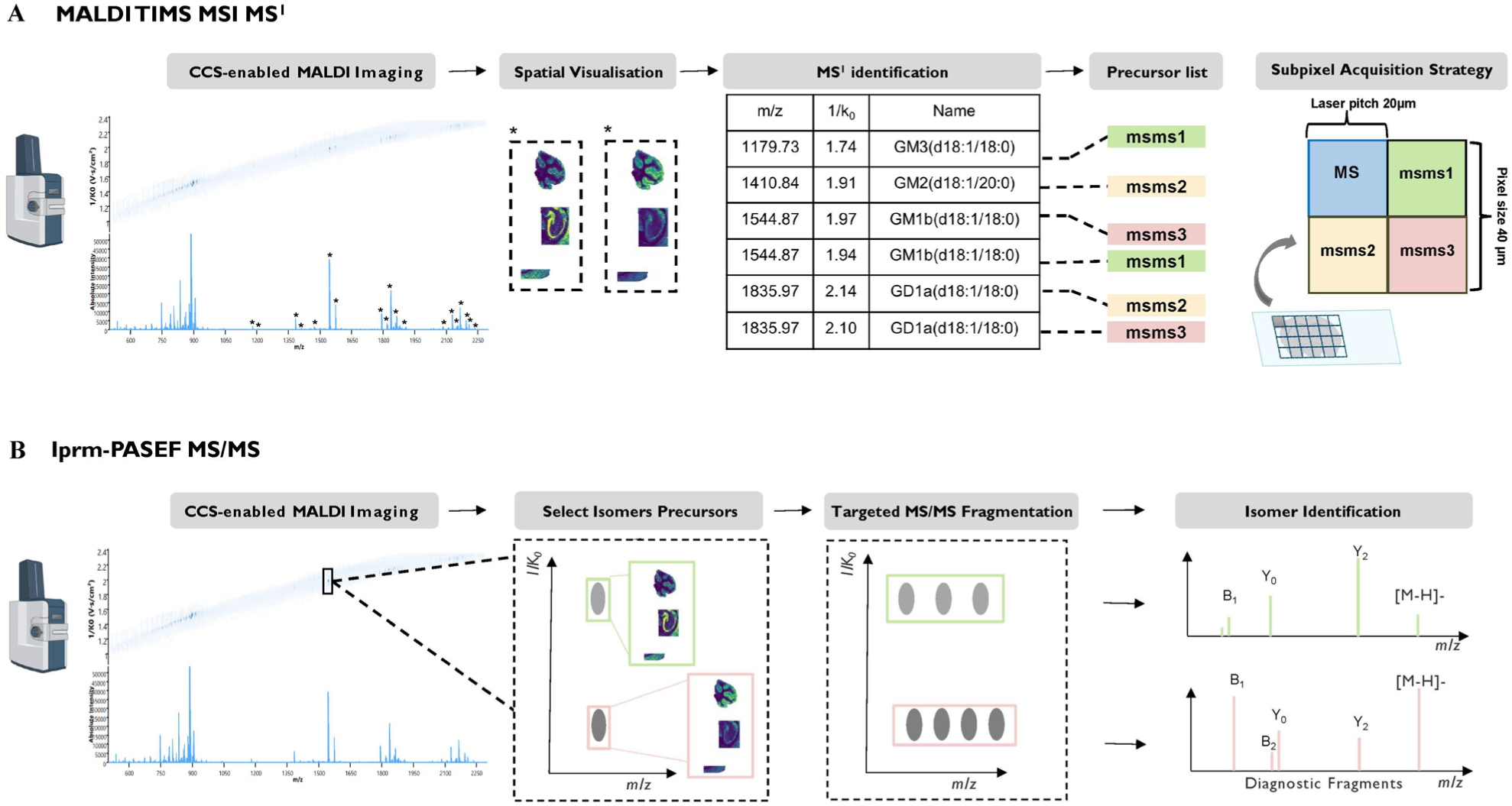
Targeted strategy for Isomer-Resolved Analysis by iPRM-PASEF-MS/MS. (A) The MALDI TIMS MS^1^ survey scan is acquired from which a precursor list is selected based on MS^1^ annotations and their mobility (1/k0) values. The MSI setup uses a 40 μm pixel size, which is subdivided into four 20μm pixels for acquisition. The first quadrant was used for MS^1^ acquisition (blue), and the remaining three quadrants are available for the following iPRM-PASEF MS/MS runs. Specifically, msms1 (green) included 21 selected precursors, msms2 (yellow) 16 precursors, and msms3 (red) 13 precursors. In total 50 precursors were analyzed with iPRM-PASEF-MS/MS (B) Selected precursor ions are isolated based on mobility (1/k0) values and CID fragmented. Gangliosides are identified based on their fragmentation patterns.

Based on the MS^1^ dataset, 50 out of 63 mobility-resolved GG features were selected for targeted MS/MS fragmentation across three iPRM-PASEF acquisition rounds (Fig. 2A,B), as explained below. In principle, each iPRM acquisition cycle allowed to target up to 25 precursor ions, provided their ion mobility profiles do not overlap. However, due to the crowded ion mobility distribution of the GG species, the final selection was constrained by partially overlapping features. Therefore, only 50 out of the 63 candidate ganglioside ion species were included for MS/MS fragmentation analysis. We prioritized detailed MS/MS data interpretation of ganglioside species that displayed distinct spatial distributions between *Npc2+/+, Npc2-/-,* and AAV-BR1-NPC2 treated *Npc2-/- vehicle* mice, as these species were considered most relevant for investigating disease-associated alterations and therapeutic responses.

These analyses enabled the structural characterization of mobility-resolved GG features that could not be distinguished based on MS^1^ data alone. In the following sections, we focus on selected ganglioside species to demonstrate the ability of the workflow to resolve isomeric structures directly in tissue while preserving spatial information.

### MS^1^-detection and spatial distribution of simple gangliosides

Neuronal cells are particularly enriched in acidic GSLs, including gangliosides, which explains why defects in GG degradation have detrimental impact on the nervous system [25, 26]. Lipid profiling by MALDI MSI of *Npc2-/-* mouse brains revealed a broad mass range of gangliosides across brain tissue compartments. Confident annotations based on exact mass alone were limited to simpler GG species, including GM3, GM2, GD3, and GD2 species (Fig. 3A). These ions exhibited a single, well-defined mobility feature per *m/z* value, consistent with a simple oligosaccharide composition and the absence of structural isomeric overlap. Representative species include GM3(d18:1/18:0) at *m/z* 1179, GM2 d18:1/18:0 at *m/z* 1382, and GM2(d18:1/20:0) at *m/z* 1410, reflecting the diversity in the ceramide backbone of GG, and a predominance of d18:1 species in neuronal tissue (Fig. 3B,C and Fig. S3) [27, 28].

**Figure 3:**
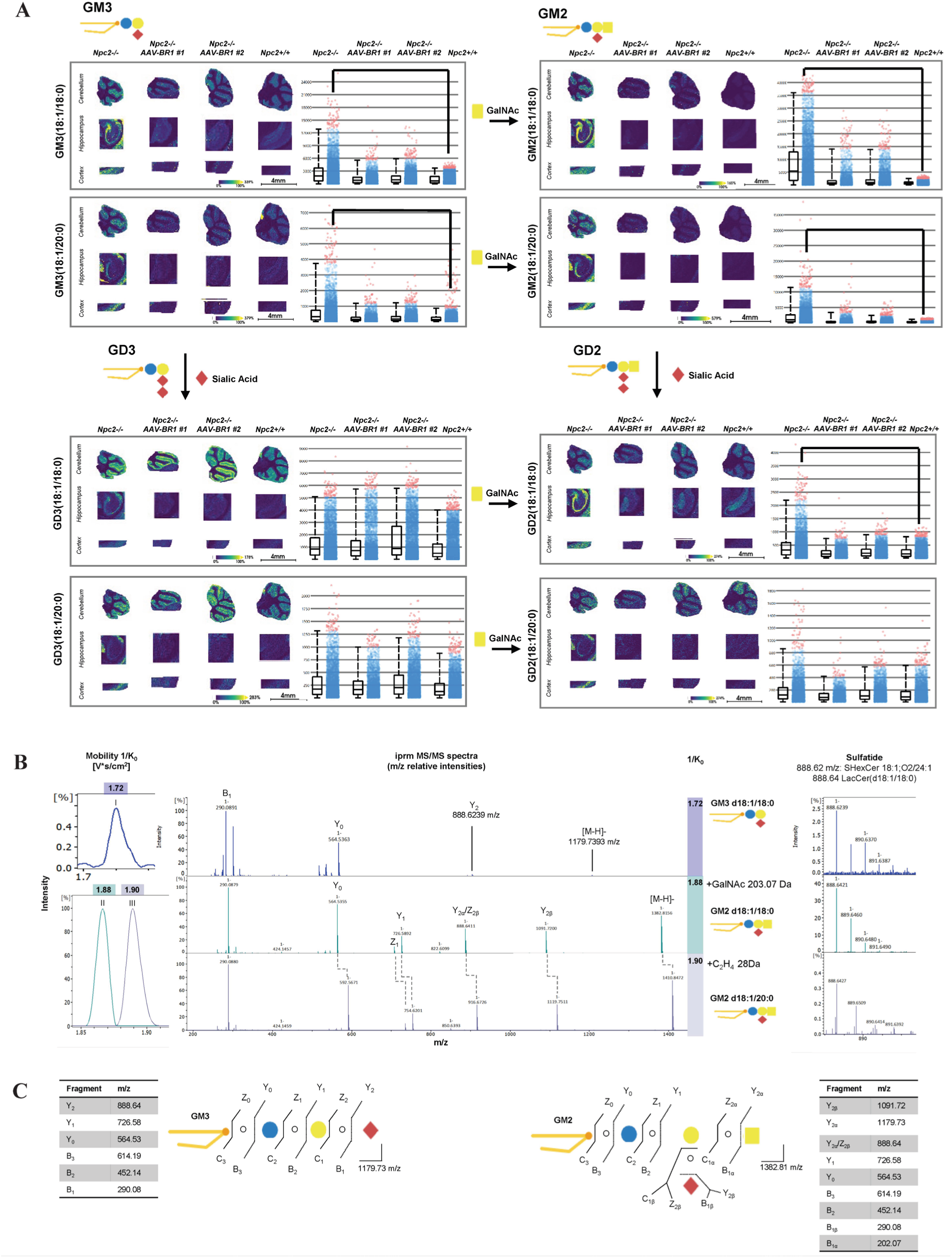
Spatial mapping and iPRM-PASEF-MS/MS analysis of simple ganglioside species in mouse brain. A) MSI showing the spatial distribution of simple gangliosides GM3, GM2, GD3, and GD2, in brain sections. Each GG is represented by varying ceramide composition (d18:1/18:0 or d18:1/20:0). The adjacent boxplot displays RMS-normalized ion intensities across samples, showing the relative abundances of the GGs between the phenotypes. B) iPRM-PASEF-MS/MS spectra of selected GGs. Top GM3(d18:1/18:0) in purple; middle, GM2(d18:1/18:1), with a mass increase of 203 Da of GalNAc relative to GM3; bottom, GM2(d18:1/20:0) in grey, showing an additional 28 Da mass difference corresponding to the longer ceramide chain. C) Structural representation of GM3 and GM2, indicating oligosaccharide composition. Fragment ion nomenclature (Y, B; Z, C) is annotated, indicating the diagnostic fragments used for MS/MS-based identification.

Spatially, these GGs were detected in the hippocampus, cortex, and cerebellum, primarily accumulating in small, spherical structures (Fig. 3A). The 20 μm spatial resolution was insufficient to resolve individual lysosomal compartments, which range from 0.1 μm to 1.2 μm. However, higher-resolution MSI at 5 µm provided additional detail, confirming the enrichment of GM3 and GM2 in these regions and revealing close spatial proximity with partial overlap (Fig. 4). Although histological validation of the observed spatial patterns was not possible on the same tissue section after MALDI-2 imaging due to tissue depletion during acquisition, the observed GG distributions are consistent with previous reports of ganglioside accumulation in NPC models [14].

**Figure 4:**
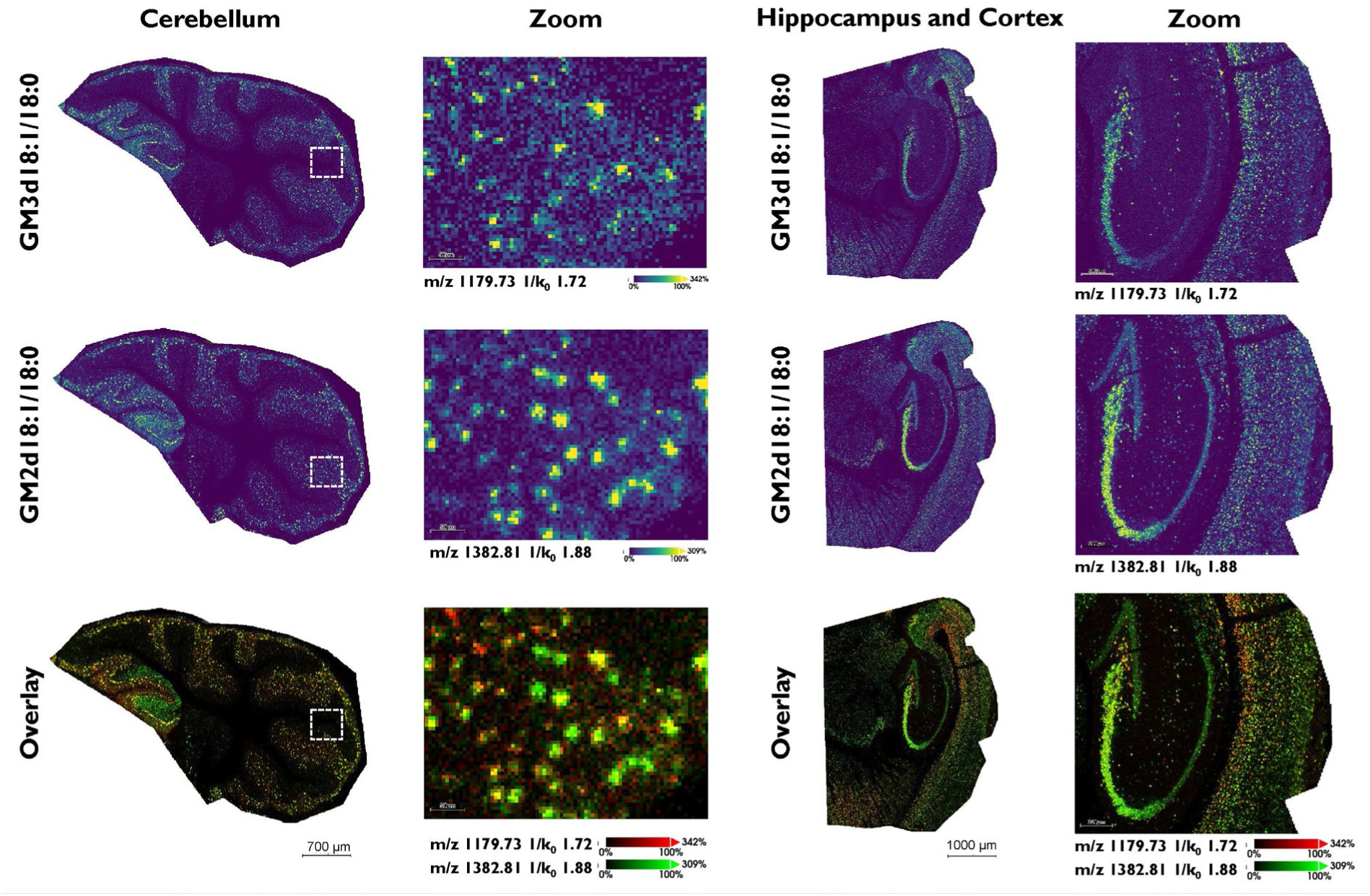
Co-localization of GM3 and GM2 revealed by 5µm microGRID MS imaging. 1179.73 m/z 1/k0 1.72, and GM2(d18:1/18:0): 1382.81 m/z 1/k0 1.88, in the cerebellum and hippocampus of a Npc2-/- mouse. The first two rows show the spatial distributions of GM3 and GM2 intensities, while the bottom row presents an overlay of both ions: red-channel GM3 (m/z 1179.73) and green-channel GM2 (m/z 1382.81), illustrating their localization and partial overlap within tissue.

Multiple mechanisms may contribute to the accumulation of GM2 and GM3 observed in *Npc1-/-* models. One proposed mechanism suggests that accumulating cholesterol inhibits the recruitment or activity of sphingolipid binding proteins, such as saponins (Sap A, Sap B), or the GM2 activator protein, to intra-luminal vesicles, which drives the lysosomal degradation of complex GGs [29]. Reduced function of these proteins impairs substrate presentation to hydrolases and promotes the buildup of GM2, GM3, and lactosylceramides. Consistent with this mechanism, *Npc2-/-* mice display ganglioside abnormalities similar to those described in Npc1-/- mice. In summary, clear spatial patterns and confident structural assignments can be established for simpler GGs, providing a foundation for exploring more complex isomers, thereby enabling an in-depth analysis of the metabolic turnover of complex GGs.

### Structural validation of simple gangliosides and co-isolation effects

To confirm the identity of the low-mass GGs detected at the MS^1^-level, targeted iPRM-MS/MS was performed directly on tissue of ion signals at *m/z* 1179.73 and 1382.81. Fragmentation patterns validated the expected ceramide backbones (d18:1/18:0) and (d18:1/20:0) and confirmed the structural assignments of GM2 and GM3 (Fig. 3B).

A sulfatide species, SHexCer(18:1;O2/24:1) at *m/z* 888.62, was also detected in the MS/MS dataset (Fig. S3). A low-abundance ion corresponding to this sulfatide was observed in the iPRM-fragmentation spectrum of the isolated GM3 precursor at *m/z* 1179.73 (Fig. 3B). In the MS^1^ images, SHexCer showed high intensity in distinct regions that did not spatially overlap with GM3(d18:1/18:0). On-tissue MS/MS fragmentation of the GM3 precursor yielded the expected LacCer(d18:1/18:0) backbone at *m/z* 888.64, confirming the presence of GM3(d18:1/18:0). Additional fragmentation of the isolated SHexCer species at *m/z* 888.62 produced a distinct spectrum with diagnostic fragment ions not observed in the GM3 iPRM-PASEF-MS/MS spectrum, confirming that these signals originate from different lipid classes (Supplementary Figure 3). However, the detection of the *m/z* 888.62 fragment in the GM3 iPRM-PASEF-MS/MS spectrum indicates that an unknown species different from the expected LacCer backbone is present (Fig. 3B). It remains unresolved whether this signal derives from a minor sulfated variant of GM3 or from the co-isolation of an unknown species within the precursor window [30]. However, no other candidates were observed within the mobility or mass spectrum window that could explain this co-isolation. The SHexCer sulfatide exhibits a distinct mobility value of 1/K_0_ 1.50 compared to GM3, with 1/K_0_ 1.72, indicating that TIMS separation clearly resolves these species at the MS^1^-level.

### Resolved GM1 isomers reveal structural and spatial differences in mouse brain

In addition to differentiating the simpler GGs, larger, more complex GGs were also detected and resolved within the mobility window. At *m/z* 1544.87, corresponding to GM1 isomers, three distinct mobilities were observed with 1/K_0_ of 1.92, 1.94, and 1.97, respectively. The iPRM-PASEF-MS/MS spectra yielded isomer-specific fragmentation behavior, enabling differentiation and assignment of 1/K_0_ 1.92 and 1.97 to GM1b and GM1a, respectively (Fig 5A). In particular, 1/K_0_ 1.92 displayed fragment ions consistent with GM1b, including the B_2_ fragment NeuAc-GalNAc-Glc (*m/z* 655). In contrast, the 1/K_0_ 1.94 and 1/K_0_ 1.97 mobilities exhibited the characteristic neutral loss of the B_2α_ GalNAc-Glc fragment (365 Da), consistent with GM1a (Fig. 5B,C). These GG isomer assignments are consistent with previous reports that branched structures like GM1a adopt a more bulky three-dimensional conformation in the gas phase relative to linear gangliosides like GM1b, resulting in a measurable shift in mobilities at high TIMS resolving power [11].

**Figure 5:**
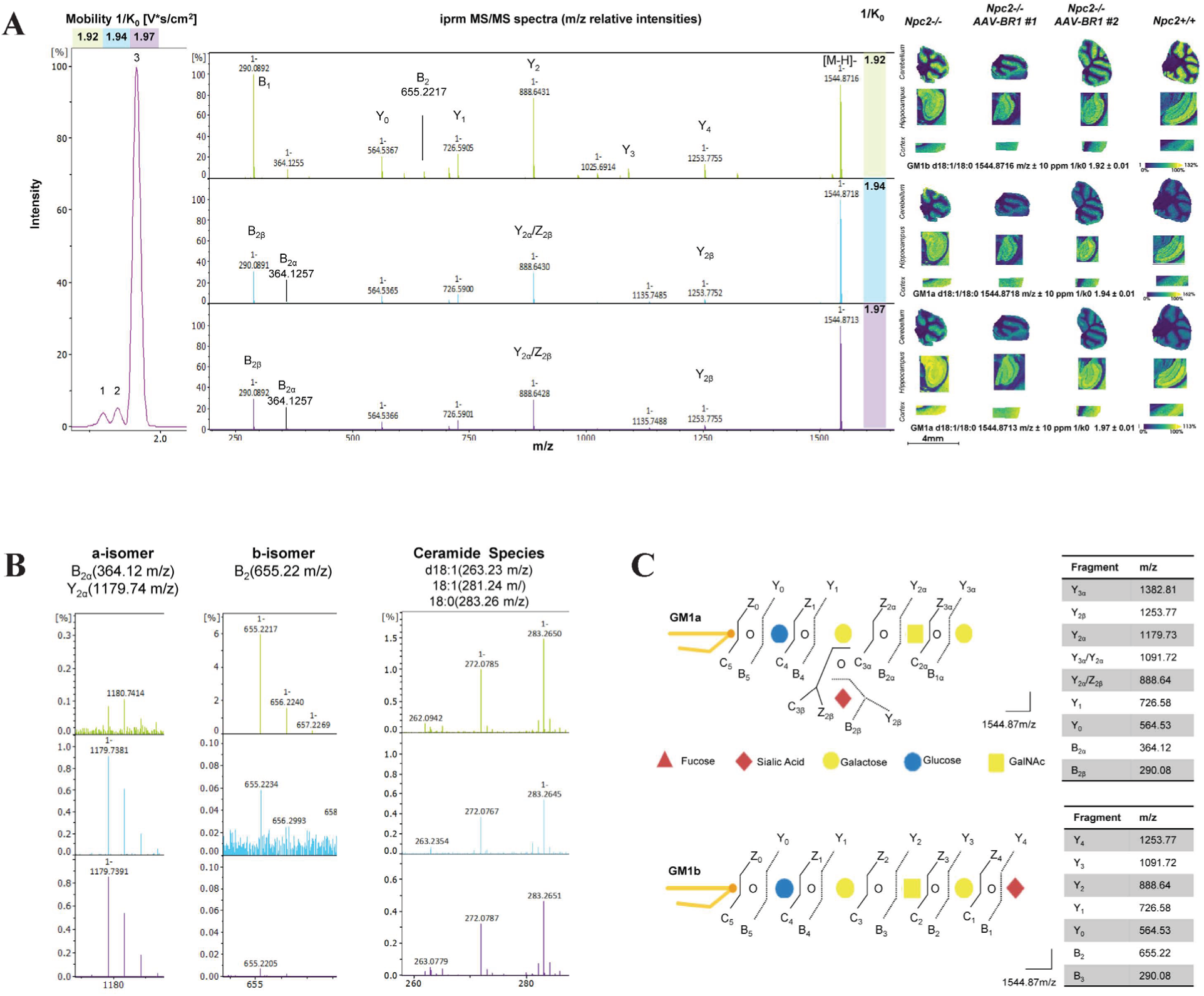
iPRM-PASEF-MS/MS and spatial mapping of GM1 isomers in mouse brains. A) iPRM-PASEF-MS/MS spectra of GM1 isomers detected at 1/k0 1.92, 1.94, and 1.97. Adjacent MALDI MSI show the GM1 distributions in Npc2-/-, Npc2-/- AAV-BR1-NPC2 (#1 and #2), and Npc2+/+ mice at 20µm resolution. B) Zoom-in of unique fragments to GM1a, GM1b, and masses corresponding to different ceramide tail composition. C) Structural representation of GM1a and GM1b, indicating the oligosaccharide composition. Fragment nomenclature (Y, B; Z, C) is shown, with a table summarizing the unique fragment ions used for isomer identification.

Interestingly, each mobility-resolved MS/MS spectrum contained fragment ions corresponding to carbon chain saturations of: d18:1, d18:0, 18:1, and 18:0 (Fig. 5B). This suggests that positional differences in ceramide saturation do not produce sufficient topological changes to allow separation by ion mobility by current TIMS technology. Accordingly, the observed isomer separation reflects differences in the GG oligosaccharide composition rather than variation in the ceramide component.

Across the brain tissue, GM1a displayed higher intensity than GM1b, consistent with its well-established abundance in neuronal membranes [31]. Within the cerebellum, the 1/k_0_ 1.94 and 1.97 were moderately elevated in *Npc2-/-* relative to *Npc2+/+*, suggesting that even modest shifts in GM1a distribution occur in NPC2-deficiency (Fig. 5A). The magnitude is lower than that observed for simple structured GGs but may still reflect altered trafficking or turnover of the GM1 species in NPC2 deficiency [32]. Interpretation should be made with caution, as subtle differences may also arise from variation in tissue section depth and the inherent heterogeneity between brain sections.

In summary, GM1 isomers are separated by TIMS based on oligosaccharide structure rather than ceramide composition, enabling their identification by TIMS-MS/MS and revealing subtle spatial distribution and abundance differences in the NPC2-deficient brain tissue.

### Fucosylated GM1 is associated with NPC2-deficiency

A fucosylated GM1a species was detected at *m/z* 1690.92 (Fig 6A). The detection of two unique fragments supported this structural assignment: B_3α_, Fuc-Gal-GalNAc at *m/z* 307.10 and Y_2α_, NeuAc-Gal-Glc-Cer at *m/z* 1179.73, indicating that the fucose residue is attached to the terminal end of the oligosaccharide chain (Fig. 6B, C). The annotated fucosylated GM1a (Fuc-GM1a) (d18:1/18:0) was detected exclusively in grey matter regions of *Npc2-/-* and AAV-BR1-NPC2-treated *Npc2-/-* mice, suggesting that alterations in fucosylation are associated with disease progression and are not fully restored by gene therapy (Fig 6D). Fucosylation of GGs is a well-known modification described in relation to cancer and inflammation, where it can influence immune signaling, cell-cell interactions, and CNS glial responses [33]. However, current literature lacks detailed descriptions of fucosylated GGs in NPC disease or other LSDs. We speculate that altered immune microenvironments and impaired GG trafficking are plausible contributors to the presence of Fuc-GM1a in NPC2-deficiency [1]. Alternatively, a lack of functional NPC2 protein may reduce fucosidase enzyme activities, resulting in increased fucosylation of glycolipids and glycoproteins, a hallmark of another LSD, fucosidosis [34]. While the present data cannot distinguish between these possibilities, the observations emphasizes a previously overlooked modification associated with disease-related changes in NPC2-deficient animals [29].

**Figure 6:**
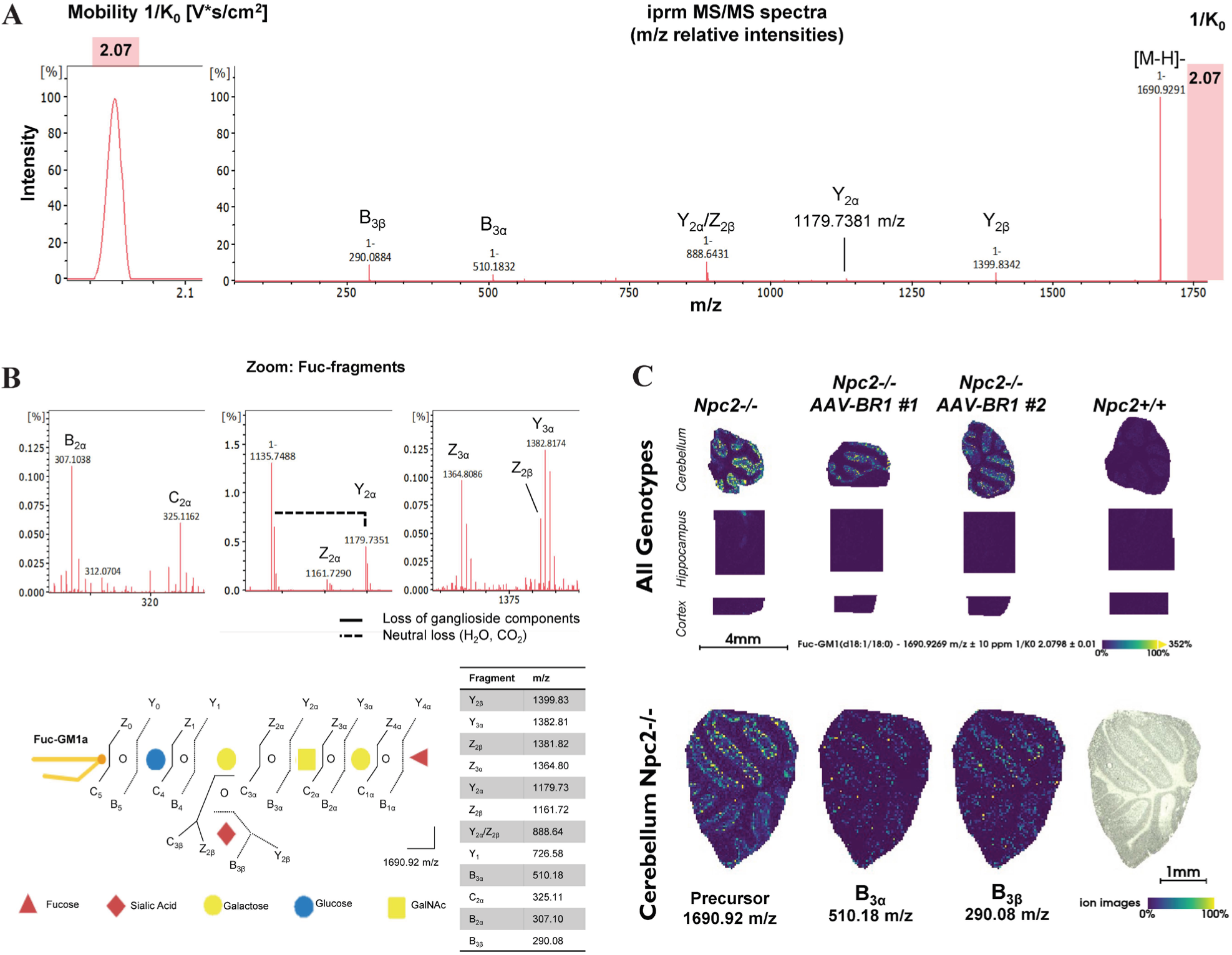
Identification of Fuc-GM1a in Npc2-/- tissue. (A) Full-range iPRM-PASEF-MS/MS spectrum of the isolated ganglioside species at m/z 1690.92. (B) zoom-in view highlighting lower-abundant but unique diagnostic fucose-containing fragments. All fragments are designated according to the established ganglioside nomenclature. (C)Summarized fragmentation pattern of Fuc-GM1a. (D) MALDI-MSI ion images of Fuc-GM1a showing spatial localization enriched within the cerebellar grey matter in Npc2-/-, Npc2-/- AAV-BR1-NPC2, and Npc2+/+ mice.

### Identification of GD1a and GD1b isomers in mouse brain

Gangliosides contain a conserved tetrasaccharide core structure that can be further modified by the addition of sialic acid residues. Gangliosides containing two sialic acids are classified as GD1 gangliosides. The different possible positions of sialic acid attachment within the oligosaccharide chain give rise to multiple GD1 structural isomers, including GD1a and GD1b.

Previous TIMS-MSI studies demonstrated the ability to separate GD1 isomers in murine kidney and brain tissue [11, 16]. Building on these observations, we investigated whether the mobility-separated features detected at m/z 1835.96 represented distinct GD1 isomeric species. Four distinct ion mobility features were detected at *m/z* 1835.96, with 1/K_0_ values of 2.08, 2.10, 2.13, and 2.15, indicative of multiple GD1 isomers (Fig. 7). The position of the sialic acids within GD1a and GD1b results in characteristic differences in neutral loss behavior that can support isomer assignment. GD1a, containing one terminal and one internal sialic acid, predominantly shows CO_2_ loss, attributed to the terminal sialic acid residue where the carboxyl group (COOH) readily fragments under MALDI conditions [35]. In contrast, GD1b contains a disialic acid arrangement in which both sialic acids are located internally, and more frequently displays neutral loss of water (H₂O). All four mobility-isolated MS/MS spectra showed CO_2_ and H_2_O losses; specifically, the 1/K_0_ 2.13 exhibited pronounced CO_2_ loss, identifying it as GD1a (Fig. 7A).

**Figure 7:**
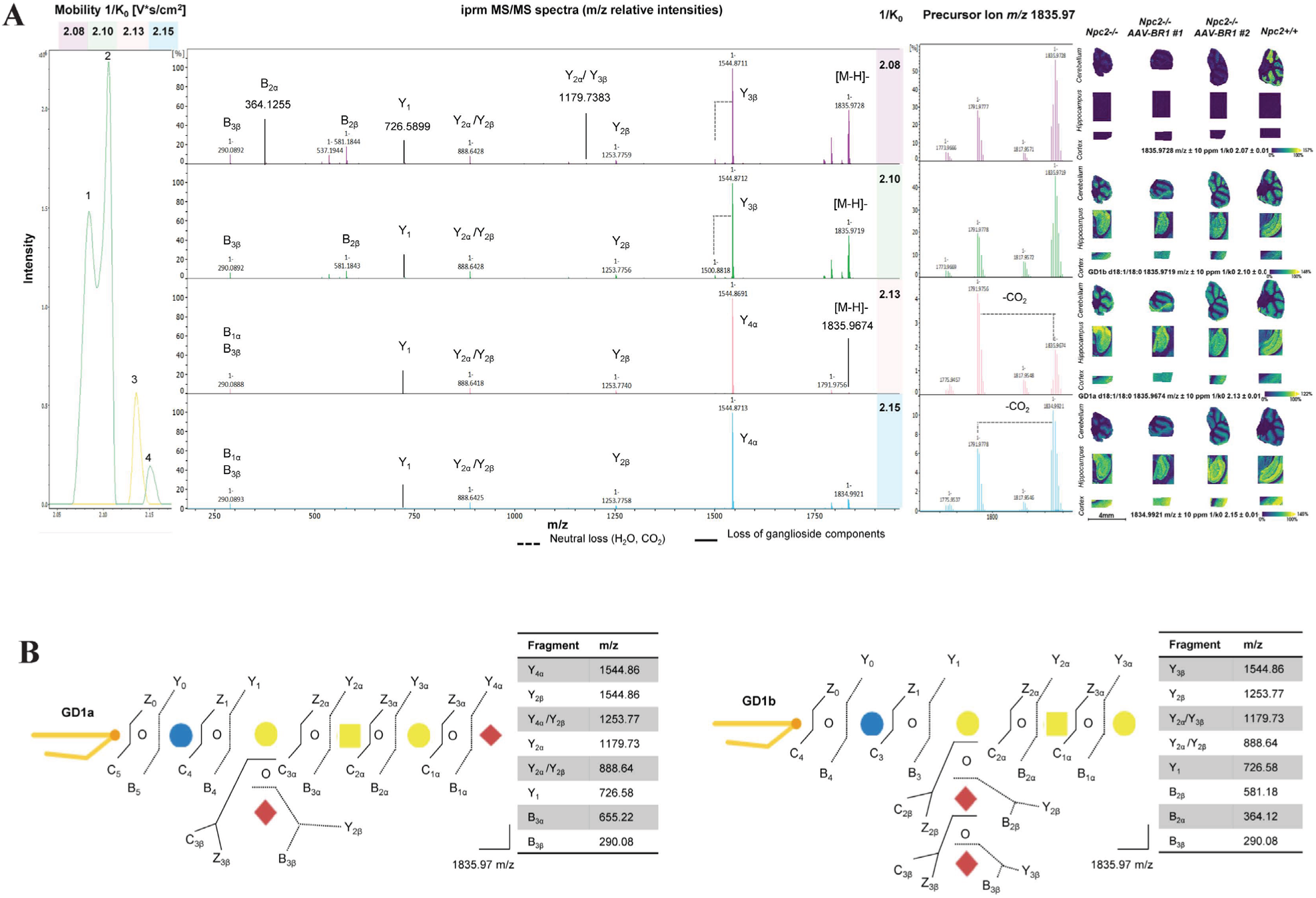
iPRM-PASEF-MS/MS differentiation and structural assignment of GD1a and GD1b isomers. A) iPRM-PASEF-MS/MS spectra acquired for precursor ion m/z 1835.96 revealed four distinct mobility features (1/k0 = 2.08, 2.10, 2.13, 2.15 V*s/cm^2^). Each feature produced unique GL fragment ion patterns annotated with Y, B, Z, and C, respectively. Neutral losses: H2O, CO2, are indicated. MALDI MSIs show distinct regional distributions of GD1 isomers across the phenotypes: Npc2-/-, Npc2-/- AAV-BR1, and Npc2+/+. B) Proposed fragmentation schemes for GD1a and GD1b isomers annotated with diagnostic Y, B, Z, and C ions. Fragment ions detected supporting structural assignments are summarized in the accompanying table.

In addition to neutral losses, the unique fragment ions observed in the iPRM-PASEF-MS/MS spectra provide further support for the assignment of the detected mobilities to GD1a or GD1b. The B_3α_ ion at *m/z* 655 (NeuAc-Gal-GalNAc) and the 1.4A cross-ring fragment at *m/z* 380.11 were detected in the specific mobility of 1/K_0_ 2.13, indicative of GD1a-type structures. In contrast, fragments corresponding to the B_2α_ ion (Gal-GalNAc, *m/z* 364.12) and the Y_2α_ ion (NeuAc-Gal-Glc-Cer(d18:1/18:0), *m/z* 1179.73) were observed in the 1/k_0_ 2.08 and 2.10 mobilities, respectively, indicative of GD1b-type structures. The disialic fragment at *m/z* 581, diagnostic of GD1b, was also present in these mobilograms (Fig. 7B). Taken together, the combination of neutral losses and unique fragment ions provides strong indications for assigning the detected mobilities of 1/K_0_ 2.13 to GD1a and 1/K_0_ 2.08, 2.10 to GD1b.

Some GD1b-associated fragments were also observed in the MS/MS spectrum of the 1/K_0_ 2.13 feature, despite the predominant presence of GD1a-associated fragments (Fig. 7A). This is likely explained by partial co-isolation of neighboring mobility species, as the GD1 features at *m/z* 1835.96 occur within closely overlapping mobility ranges. Further improvement of TIMS resolving power could reduce this effect, although at the cost of increased acquisition time.

To summarize, GD1 isomers can be differentiated by combining mobility separation with diagnostic neutral losses and fragment ions, enabling structural assignment despite partial co-isolation arising from closely overlapping mobility features.

### Altered ganglioside GD1 processing in NPC2 deficiency

The O-Ac GD1 species (*m/z* 1877.98; 1/K_0_ 2.10) was detected exclusively in the cerebellum of the *Npc2+/+* mice (Fig 8). This assignment is supported by observation of the B_2β_ fragment ion at *m/z* 623.19, consistent with a disialic acid structure carrying a single O-Ac moiety, and the B_3β_ fragment ion at *m/z* 332.09, confirming the presence of an acetylated sialic acid. Furthermore, the fragment ion at *m/z* 1544.87 corresponds to GM1a(d18:1/18:0), indicating assignment of the O-Ac moiety to the outer sialic acid of the GD1b structure (Fig. 8A, B).

**Figure 8:**
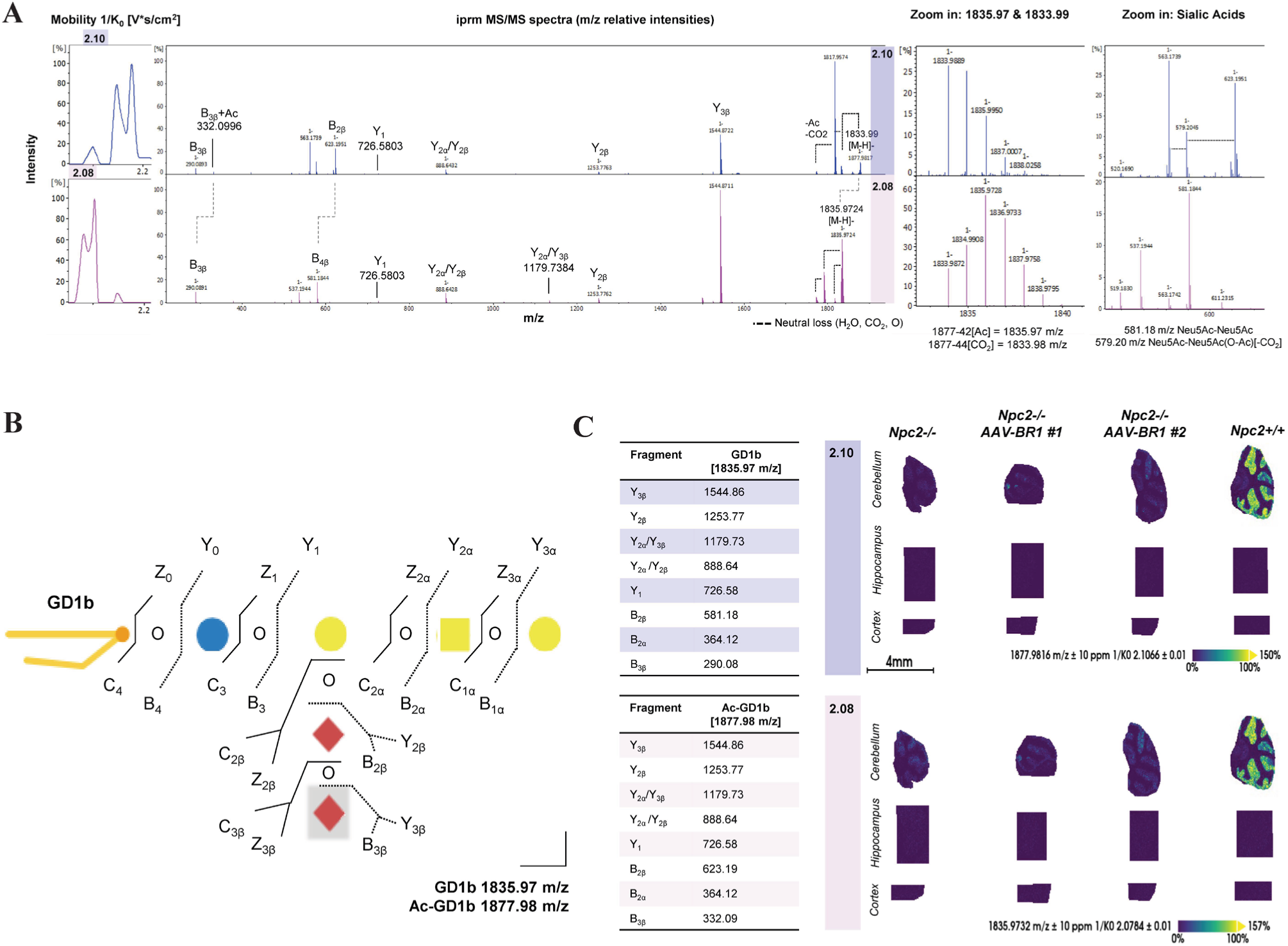
MALDI iPRM-PASEF MS/MS analysis of GD1b and O-acetylated GD1b in Npc2+/+ mouse cerebellum. A) MS/MS spectra of GD1b m/z 1835.97, 1/k0 20.8 (pink) and O-acetylated GD1b m/z 1877.98, 1/k0 (purple) in Npc2+/+. Key fragments used for identification are indicated; neutral losses (H2O, CO2, O-Ac) are shown as dashed lines. B) GD1b structure with CID fragmentation map; grey boxes indicate the proposed O-acetylation site. Tables list fragment masses for confident isomer assignment. C) MALDI-MSI showing GD1b and Ac-GD1b spatial distribution localized to the cerebellum in Npc2+/+ mouse brain.

We note that the spatial distribution of the O-Ac GD1 species overlaps with that of the non-acetylated GD1 ion species (*m/z* 1835.96, 1/K_0_ 2.08) (Fig. 8B). Labile sialic acid linkages may break during MALDI ionization, leading to in-source decay of acetylated sialic acid, thereby generating the (*m/z* 1835.96, 1/K_0_ 2.08) species from the O-Ac GD1 (*m/z* 1877.98) precursor ion [35]. The identical spatial distribution supports this interpretation (Fig. 8C).

Sialic acid O-acetylation is an enzymatically regulated modification that occurs in the Golgi during ganglioside biosynthesis [29]. Although O-Ac GGs remain poorly characterized in NPC pathology, previous studies have identified O-Ac GD3 species in brain tissue, albeit at low abundance [36]. In NPC disease, lysosomal trapping of gangliosides may reduce their accessibility to Golgi-associated modifying enzymes, providing a possible explanation for the absence of O-Ac GD1 in Npc2-/- mice (Fig. 8C). Although NPC2 gene therapy reduced neuronal cholesterol accumulation, O-acetylation remained absent in AAV-BR1-NPC2-treated mice, suggesting that ganglioside processing or trafficking was not fully restored [1]. Whether this reflects incomplete rescue, delayed metabolic recovery, or cell-type specific differences in therapeutic response remains to be resolved.

O-acetylation can occur at several positions on the sialic acid residue, potentially influencing GG interactions and biological properties, although the effects of individual modifications remain incompletely understood [37, 38]. In the present study, the analytical approach enables detection of O-Ac but does not allow assignment of the specific acetylation site. Although MS/MS fragmentation confirms the presence of an acetylated sialic acid, the acquired fragment ions do not provide sufficient structural information to distinguish between possible O-Ac positions on the sialic acid residue. Further method development would therefore be relevant to determine whether specific acetylation patterns are altered in NPC deficiency and to assess their biological influence [37].

In summary, O-Ac GD1 is detected exclusively in *Npc2+/+* tissue and absent in NPC2-deficiency, suggesting impaired ganglioside trafficking and Golgi-dependent modifications, while partially overlapping signals indicate potential in-source fragmentation of acetylated species.

## Conclusion

This study demonstrates that ganglioside alterations in NPC2 deficiency extend beyond changes in overall abundance, revealing a layer of molecular complexity defined by isomer composition, structural modifications, and spatial organization. By integrating TIMS-based ion mobility separation with spatially resolved MS/MS, we demonstrate in situ structural characterization of ganglioside species that cannot be distinguished based on mass alone. This added structural dimension provides new insight into how individual ganglioside species are affected by lysosomal dysfunction and therapeutic intervention, highlighting the importance of molecular specificity when investigating lipid alterations in disease. The persistence of specific ganglioside alterations following therapeutic intervention further emphasizes that changes in overall ganglioside abundance alone do not fully describe the molecular complexity of NPC pathology. More broadly, this approach enables spatially resolved investigation of lipid structural diversity in biological systems where molecular composition alone is insufficient to explain disease-associated changes.

## Acknowledgements

This research project was supported by a research project grant (NNF22OC0080228) and a research infrastructure grant (INTEGRA, NNF20OC0061575) from the Novo Nordisk Foundation to ONJ. DW acknowledges funding by the Danish Research Council (2034-00136B) and by the Ara Parseghian Medical Research Fund. CR and AB acknowledge the financial support provided by Direktør Emil C. Hertz og hustru Inger Hertz’ Fond, Dagmar Marshalls Fond, Hørslev-Fonden, Læge Sophus Carl Emil Friis og Hustru Olga Doris Friis Legat, and the Research Initiative for Brain Barriers and Drug Delivery (RIBBDD). The authors also thank Karina Lassen Holm and Dorte Hermansen, Aarhus University, Denmark, for their excellent technical assistance throughout the mouse study, and Associate Professor Maj Schneider Thomsen for her valuable assistance with tissue collection. Finally, Eva Hede, Aalborg University, Denmark, and Jakob Körbelin, Universitätsklinikum Hamburg-Eppendorf, Germany, are acknowledged for the design and production of the AAV-BR1-NPC2 viral vector.

