## Supplementary material for "Ganglioside profiling by ion mobility and mass spectrometry imaging identifies distinct biomolecular isomers in Niemann-Pick Disease": Suppl.Information

### Supplementary Figures

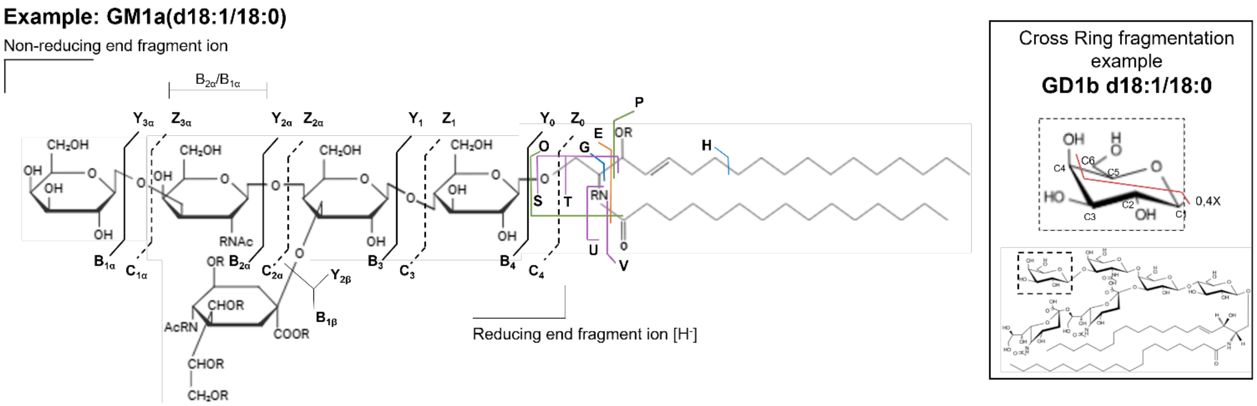


***Supplementary Figure S1: Fragmentation overview and nomenclature for gangliosides.*** *GM1a is shown as an example to illustrate how the molecule fragments during MS/MS, while a glucose residue from GD1b is used to show a typical cross-ring fragment. Fragment ions are annotated according to standard nomenclature, where B-, C-, Y-, and Z-type ions originate from glycosidic bond cleavages on the reducing and non-reducing sides of the oligosaccharide chain, respectively. Cross-ring fragments (A- and X-type) provide additional linkage information. The colour code highlights fragments derived ion from different location in the ceramide chain structure: O and P ions (green) arise from the spingoid base, S-V ions (purple) to the fatty acid chain, G and H ions (blue) to internal breaks within the spingoid base, and the E ion (orange) to fragments containing portion of both the sphingoid base and fatty acid chain.*

***Supplementary Table S1: Annotated gangliosides based on MS[1] profiles.*** *MSI data were imported into SCiLS lab, from which m/z and mobilities were manually selected in the sum mobility heatmap of the measured sample regions. The features were annotated against a LipidMaps library with search parameters: mass error of ± 10ppm, CCS deviation maximum of 4%. CCS predication was enabled.*

| m/z | 1/K0 [V·s/cm$^2$] | CCS [Å$^2$] | CCS width [±] | Name | Formula | Ion Notation |
|---|---|---|---|---|---|---|
| 1179.738 | 1.745 | 353.5539 | 1.0131 | GM3(d18:1/18:0) | C59H108N2O21 | [M-H]- |
| 1207.771 | 1.7702 | 358.5675 | 1.0128 | GM3(d18:1/20:0) | C61H112N2O21 | [M-H]- |
| 1382.821 | 1.8874 | 381.7662 | 1.0113 | GM2(d18:1/18:0) | C67H121N3O26 | [M-H]- |
| 1410.85 | 1.9115 | 386.5488 | 1.0111 | GM2(d18:1/20:0) | C69H125N3O26 | [M-H]- |
| 1470.836 | 1.9051 | 385.1133 | 1.0107 | GD3(d18:1/18:0) | C70H125N3O29 | [M-H]- |
| 1544.87 | 1.9475 | 393.5142 | 1.0103 | GM1(d18:1/18:0) | C73H131N3O31 | [M-H]- |
| 1544.87 | 1.9689 | 397.8388 | 1.0103 | GM1(d18:1/18:0) | C73H131N3O31 | [M-H]- |
| 1544.873 | 1.9947 | 403.0444 | 1.0103 | GM1(d18:1/18:0) | C73H131N3O31 | [M-H]- |
| 1572.902 | 1.9683 | 397.6417 | 1.0101 | GM1(d18:1/20:0) | C75H135N3O31 | [M-H]- |
| 1572.904 | 2.0161 | 407.3048 | 1.0101 | GM1(d18:1/20:0) | C75H135N3O31 | [M-H]- |
| 1572.905 | 1.9922 | 402.4733 | 1.0101 | GM1(d18:1/20:0) | C75H135N3O31 | [M-H]- |
| 1600.933 | 1.9902 | 402.0105 | 1.01 | GM1(d18:1/22:0) | C77H139N3O31 | [M-H]- |
| 1600.934 | 2.0273 | 409.4917 | 1.01 | GM1(d18:1/22:0) | C77H139N3O31 | [M-H]- |
| 1614.952 | 2.0308 | 410.1851 | 1.0099 | KDN-GM1(d18:1/26:1(17Z)) | C79H142N2O31 | [M]- |
| 1690.93 | 2.0779 | 419.5264 | 1.0095 | Fuc-GM1(d18:1/18:0) | C79H141N3O35 | [M-H]- |
| 1791.978 | 2.1168 | 427.19 | 1.009 | GD1b(d18:1/18:0) | C84H148N4O39 | [M-CO2-H]- |
| 1791.979 | 2.0917 | 422.1233 | 1.009 | GD1b(d18:1/18:0) | C84H148N4O39 | [M-CO2-H]- |
| 1791.98 | 2.0749 | 418.7417 | 1.009 | GD1b(d18:1/18:0) | C84H148N4O39 | [M-CO2-H]- |
| 1791.98 | 2.1396 | 431.7916 | 1.009 | GD1b(d18:1/18:0) | C84H148N4O39 | [M-CO2-H]- |
| 1817.959 | 2.1284 | 429.4737 | 1.0089 | GD1b(d18:1/18:0) | C84H148N4O39 | [M-H2O-H]- |
| 1817.96 | 2.0529 | 414.2538 | 1.0089 | GD1b(d18:1/18:0) | C84H148N4O39 | [M-H2O-H]- |
| 1817.963 | 2.0924 | 422.2244 | 1.0089 | GD1b(d18:1/18:0) | C84H148N4O39 | [M-H2O-H]- |
| 1817.966 | 2.1085 | 425.4651 | 1.0089 | GD1b(d18:1/18:0) | C84H148N4O39 | [M-H2O-H]- |
| 1835.969 | 2.0795 | 419.591 | 1.0089 | GD1b(d18:1/18:0) | C84H148N4O39 | [M-H]- |
| 1835.971 | 2.1404 | 431.8762 | 1.0089 | GD1b(d18:1/18:0) | C84H148N4O39 | [M-H]- |

| | | | | | | |
|---|---|---|---|---|---|---|
| 1835.971 | 2.1017 | 424.0521 | 1.0089 | GD1b(d18:1/18:0) | C84H148N4O39 | [M-H]- |
| 1835.973 | 2.1537 | 434.5528 | 1.0089 | GD1b(d18:1/18:0) | C84H148N4O39 | [M-H]- |
| 1845.987 | 2.1477 | 433.3247 | 1.0088 | GD1b(d18:1/20:0) | C86H152N4O39 | [M-H2O-H]- |
| 1857.953 | 2.1465 | 433.0642 | 1.0088 | GD1b(d18:1/18:0) | C84H148N4O39 | [M+Na-2H]- |
| 1857.953 | 2.1286 | 429.4589 | 1.0088 | GD1b(d18:1/18:0) | C84H148N4O39 | [M+Na-2H]- |
| 1864 | 2.1661 | 437.0099 | 1.0087 | GD1b(d18:1/20:0) | C86H152N4O39 | [M-H]- |
| 1864 | 2.1781 | 439.4244 | 1.0087 | GD1b(d18:1/20:0) | C86H152N4O39 | [M-H]- |
| 1864.005 | 2.1246 | 428.6274 | 1.0087 | GD1b(d18:1/20:0) | C86H152N4O39 | [M-H]- |
| 1864.007 | 2.1465 | 433.0464 | 1.0087 | GD1b(d18:1/20:0) | C86H152N4O39 | [M-H]- |
| 1873.918 | 2.1388 | 431.4842 | 1.0087 | GD1a (NeuAc/NeuGc)(d18:1/18:0) | C84H148N4O40 | [M+Na-2H]- |
| 1873.919 | 2.1081 | 425.2803 | 1.0087 | GD1a (NeuAc/NeuGc)(d18:1/18:0) | C84H148N4O40 | [M+Na-2H]- |
| 1873.923 | 2.1539 | 434.5224 | 1.0087 | GD1a (NeuAc/NeuGc)(d18:1/18:0) | C84H148N4O40 | [M+Na-2H]- |
| 1873.925 | 2.1711 | 437.9946 | 1.0087 | GD1a (NeuAc/NeuGc)(d18:1/18:0) | C84H148N4O40 | [M+Na-2H]- |
| 1877.978 | 2.1053 | 424.7124 | 1.0087 | GD1a (NeuGc/NeuGc)(d18:1/20:0) | C86H152N4O41 | [M-H2O-H]- |
| 1877.981 | 2.1343 | 430.5773 | 1.0087 | GD1a (NeuGc/NeuGc)(d18:1/20:0) | C86H152N4O41 | [M-H2O-H]- |
| 1877.982 | 2.16 | 435.7616 | 1.0087 | GD1a (NeuGc/NeuGc)(d18:1/20:0) | C86H152N4O41 | [M-H2O-H]- |
| 1877.983 | 2.1852 | 440.841 | 1.0087 | GD1a (NeuGc/NeuGc)(d18:1/20:0) | C86H152N4O41 | [M-H2O-H]- |
| 1878.017 | 2.2044 | 444.716 | 1.0087 | Neu5Ac)GD1a(d18:1/24:1(15Z)) | C88H155N3O39 | [M]- |
| 2039.064 | 2.2678 | 457.2427 | 1.0081 | GalNAc-GD1a(Neu5Ac/Neu5Gc)(d18:1/20:0) | C94H165N5O45 | [M-CO2-H]- |
| 2039.089 | 2.2329 | 450.1873 | 1.0081 | Fuc-GD1b(d18:1/22:0) | C94H166N4O43 | [M]- |
| 2039.09 | 2.2133 | 446.2464 | 1.0081 | Fuc-GD1b(d18:1/22:0) | C94H166N4O43 | [M]- |
| 2039.09 | 2.2568 | 455.018 | 1.0081 | Fuc-GD1b(d18:1/22:0) | C94H166N4O43 | [M]- |
| 2040.056 | 2.2672 | 457.118 | 1.0081 | GalNAc-GD1a(d18:1/18:0) | C92H161N5O44 | [M]- |
| 2065.071 | 2.2584 | 455.2988 | 1.008 | GalNAc-GD1a(Neu5Ac/Neu5Gc)(d18:1/20:0) | C94H165N5O45 | [M-H2O-H]- |
| 2083.075 | 2.2739 | 458.3908 | 1.008 | GT1c(d18:1/18:0) | C95H165N5O47 | [M-CO2-H]- |
| 2083.078 | 2.2482 | 453.2199 | 1.008 | GT1c(d18:1/18:0) | C95H165N5O47 | [M-CO2-H]- |
| 2109.055 | 2.2799 | 459.5661 | 1.0079 | GT1c(d18:1/18:0) | C95H165N5O47 | [M-H2O-H]- |
| 2127.065 | 2.291 | 461.7878 | 1.0078 | GT1c(d18:1/18:0) | C95H165N5O47 | [M-H]- |
| 2127.067 | 2.2736 | 458.2738 | 1.0078 | GT1c(d18:1/18:0) | C95H165N5O47 | [M-H]- |
| 2165.007 | 2.2689 | 457.2819 | 1.0077 | GT1b(d18:1/18:0) | C95H165N5O48 | [M+Na-2H]- |
| 2165.009 | 2.2418 | 451.8213 | 1.0077 | GT1b(d18:1/18:0) | C95H165N5O48 | [M+Na-2H]- |
| 2165.011 | 2.2978 | 463.0947 | 1.0077 | GT1b(d18:1/18:0) | C95H165N5O48 | [M+Na-2H]- |
| 2193.041 | 2.3197 | 467.4766 | 1.0076 | GT1b(d18:1/20:0) | C97H169N5O48 | [M+Na-2H]- |
| 2193.041 | 2.3079 | 465.0897 | 1.0076 | GT1b(d18:1/20:0) | C97H169N5O48 | [M+Na-2H]- |
| 2193.041 | 2.2887 | 461.225 | 1.0076 | GT1b(d18:1/20:0) | C97H169N5O48 | [M+Na-2H]- |
| 2193.041 | 2.2681 | 457.0762 | 1.0076 | GT1b(d18:1/20:0) | C97H169N5O48 | [M+Na-2H]- |
| 2207.023 | 2.331 | 469.7348 | 1.0076 | Unknown | Unknown | Unknown |
| 2207.023 | 2.3121 | 465.9188 | 1.0076 | Unknown | Unknown | Unknown |
| 2207.023 | 2.2859 | 460.6491 | 1.0076 | Unknown | Unknown | Unknown |
| 2207.023 | 2.2565 | 454.713 | 1.0076 | Unknown | Unknown | Unknown |

**A**

**msms1**

| mass [m/z] | mobility start [1/K0] | mobility end [1/K0] |
|---|---|---|
| 1179.73 | 1.74 | 1.75 |
| 1207.77 | 1.76 | 1.77 |
| 1382.82 | 1.88 | 1.89 |
| 1410.85 | 1.90 | 1.91 |
| 1544.86 | 1.94 | 1.95 |
| 1544.86 | 1.96 | 1.97 |
| 1600.93 | 1.98 | 1.99 |
| 1572.90 | 2.01 | 2.02 |
| 1817.95 | 2.04 | 2.05 |
| 1835.96 | 2.07 | 2.08 |
| 1835.97 | 2.09 | 2.10 |
| 1791.97 | 2.1138 | 2.1178 |
| 1791.98 | 2.13 | 2.14 |
| 1835.97 | 2.1517 | 2.1547 |
| 1864.00 | 2.1615 | 2.1617 |
| 1864.00 | 2.17 | 2.18 |
| 2039.09 | 2.20 | 2.21 |
| 2165.00 | 2.23 | 2.24 |
| 2039.06 | 2.26 | 2.27 |
| 2207.02 | 2.2809 | 2.2899 |
| 2193.04 | 2.31 | 2.32 |

**msms2**

| mass [m/z] | mobility start [1/K0] | mobility end [1/K0] |
|---|---|---|
| 1470.83 | 1.90 | 1.91 |
| 1572.90 | 1.96 | 1.97 |
| 1572.90 | 1.98 | 1.99 |
| 1600.93 | 2.02 | 2.03 |
| 1690.92 | 2.0729 | 2.0799 |
| 1817.96 | 2.0904 | 2.0954 |
| 1873.91 | 2.10 | 2.11 |
| 1864.00 | 2.12 | 2.12 |
| 1835.97 | 2.13 | 2.14 |
| 1873.92 | 2.1529 | 2.1589 |
| 1877.98 | 2.18 | 2.19 |
| 2039.08 | 2.22 | 2.23 |
| 2039.09 | 2.2518 | 2.2578 |
| 2193.04 | 2.28 | 2.29 |
| 2193.04 | 2.30 | 2.31 |
| 2207.02 | 2.32 | 2.33 |

**msms3**

| mass [m/z] | mobility start [1/K0] | mobility end [1/K0] |
|---|---|---|
| 1544.97 | 1.98 | 1.99 |
| 1614.95 | 2.02 | 2.03 |
| 1791.98 | 2.06 | 2.07 |
| 1791.97 | 2.08 | 2.09 |
| 1877.97 | 2.10 | 2.11 |
| 1817.95 | 2.12 | 2.13 |
| 1864.00 | 2.1415 | 2.1485 |
| 1877.98 | 2.15 | 2.16 |
| 1878.01 | 2.19 | 2.20 |
| 2065.07 | 2.25 | 2.26 |
| 2127.06 | 2.2706 | 2.2776 |
| 2127.06 | 2.2.28 | 2.29 |
| 2207.02 | 2.28 | 2.29 |

**B**

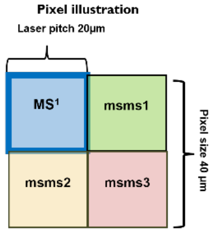


***Supplementary Figure S2: iPRM-PASEF-MS/MS precursor list and acquisition scheme.*** *A) Precursor ions were distributed across three target list (msms1:Green, msms2:Yellow, msms3:red) corresponding to iPRM-PASEF-MS/MS acquisition cycles to minimize overlap between mobility isolation windows. The table summarizes the targeted m/z values, and their respective mobility start and end positions (1/k$_0$). B) Each imaging pixel 40µm was divided into four quadrants, with one used for standard MS$^1$ acquisition (blue) and three used sequentially for targeted MS/MS measurements (msms1-3).*

*Supplementary Table S1: Spray parameters used for matrix application*

| | |
|---|---|
| Matrix | 2.5 DHAP |
| Solvent | 70% EtOH, 0.1% TFA |
| Concentration | 5mg/mL |
| Temperature Celsius | 80 |
| Pressure (psi) | 10 |
| Flow Rate (µL/min) | 100 |
| Velocity (mm/min) | 1200 |
| Track Space (mm) | 2 |
| Number of passes | 15 |
| Pattern | CC |
| Drying Time (Sec) | 10 |
| Matrix Density (mg/mm$^2$) | 3.1250 |
| Linear flow rate (mL/mm) | 8.33E-002 |

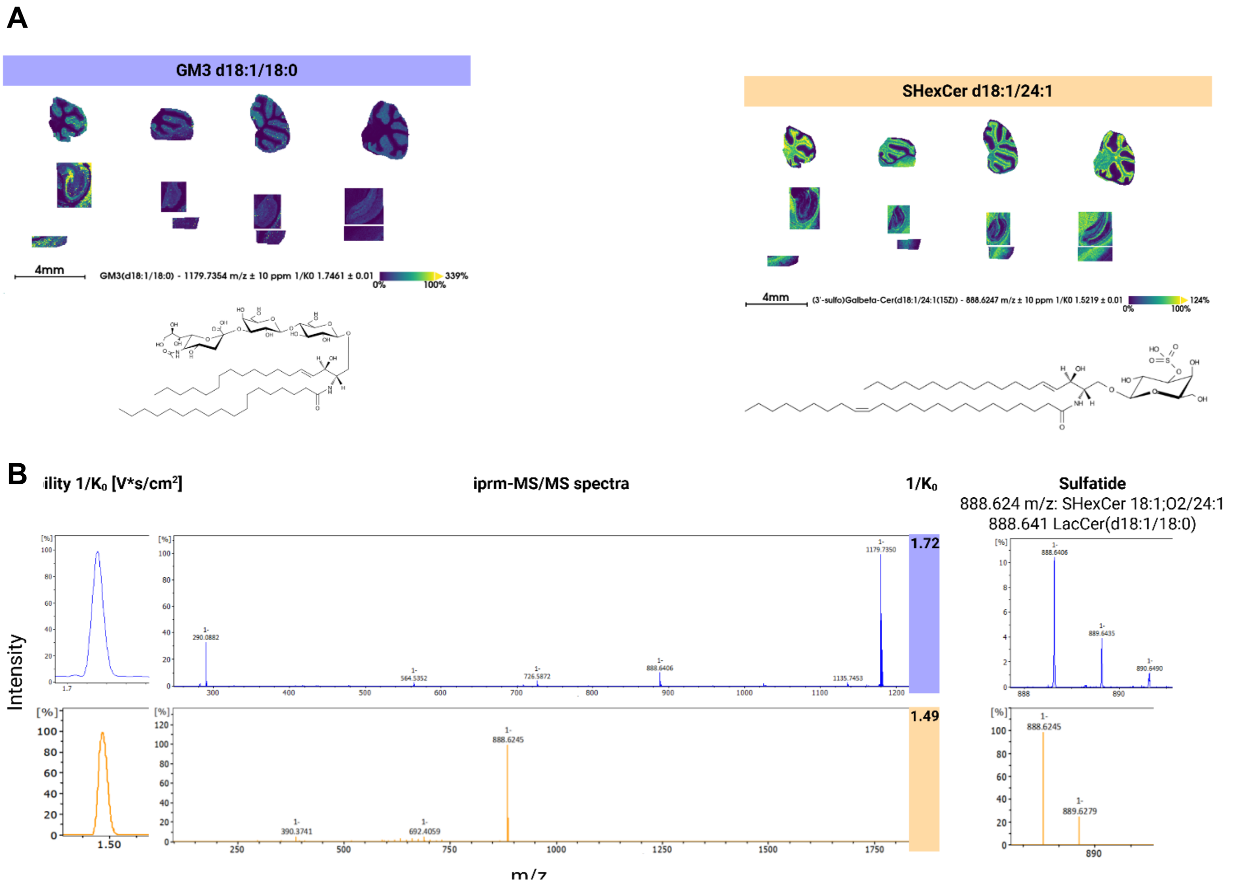


***Supplementary Figure S3: Spot MALDI confirms absence of sulfatide signal in GM3 fragmentation.*** *A) Ion images of GM3 d18:1/18:0 and SHexCer illustrate distinct spatial locations in mouse brain tissue. B) Spot fragmentation of GM3 d18:1/18:0 m/z 1179.73 produces the expected LacCer fragment without detection of the sulfatide at m/z 888.62. In contrast, SHexCer produces distinct fragment spectrum with diagnostics fragments not identified in the spot GM3 fragment spectra, nor the GM3 iPRM spectra.*
